# Frequency-dependent viscosity of salmon ovarian fluid has biophysical implications for sperm-egg interactions

**DOI:** 10.1101/2022.06.22.497200

**Authors:** Marco Graziano, Swomitra Palit, Anand Yethiraj, Simone Immler, Matthew J.G. Gage, Craig F. Purchase

## Abstract

Gamete-level sexual selection of externally fertilising species is usually achieved by modifying sperm behaviour with mechanisms thought to alter the chemical environment in which gametes perform. In fish this can be accomplished through the ovarian fluid, a substance released with the eggs at spawning. While its biochemical effects in relation to sperm energetics have been investigated, the influence of the physical environment in which sperm compete remains poorly explored. Our objective was therefore to gain insights on the physical structure of this fluid and potential impacts on reproduction. Using soft-matter physics approaches of steady-state and oscillatory viscosity measurements, we subjected salmon ovarian fluids to variable shear stresses and frequencies resembling those exerted by sperm swimming through the fluid near eggs. We show that this fluid, which in its relaxed state is a gel-like substance, displays a non-Newtonian viscoelastic and shear-thinning profile, where the viscosity decreases with increasing shear rates. We concurrently find that this fluid obeys the Cox-Merz rule below 7.6 Hz and infringes it above, thus indicating a shear-thickening phase where viscosity increases provided it is probed gently enough. This suggests the presence of a unique frequency-dependant structural network with relevant implications on sperm energetics and fertilisation dynamics.

## Introduction

The micro-conditions of fertilization are poorly understood in the majority of animal species (Cosson, 2015; Eisenbach & Giojalas, 2006; Kholodnyy et al., 2019). Following ejaculation, sperm find and fertilise eggs, but this usually takes place in the presence of post-mating sexual selection arising from sperm competition with rival males (Birkhead & Pizzari, 2002; Parker, 2020), and cryptic female choice that biases paternity (Firman et al., 2017). We now know that polyandry (female mating with multiple males in a given breeding episode) is widespread and common in nature (Taylor et al., 2014), and that post-mating sexual selection plays a crucial role in governing reproductive fitness (Simmons, 2005). It is likely to be responsible for the tremendous diversity in sperm morphology (Ramón et al., 2014; Pitnick, Hoksen and & Birkhead, 2008) and female reproductive tract morphological complexity (Kelly & Moore, 2016; Sloan & Simmons, 2019). Although many studies have revealed the importance of post-mating sexual selection for dictating variance in individual fertilization success (Gasparini & Pilastro, 2011; Kekalainen & Evans, 2018; Lüpold et al., 2012), we still understand little about the exact mechanisms that control the outcome of post-mating sexual selection and, ultimately, individual fertilisation success (Birkhead & Pizzari, 2002).

In terms of female control over paternity, internal fertilisation clearly offers greater direct opportunity to manage sperm and the fertilisation process, compared with external fertilisation. In internal fertlizers, sperm are deposited within the female reproductive tract, and then move from the insemination site either directly towards the egg for fertilisation, or indirectly via short-or long-term storage. Sperm can move under their own flagellar propulsion, or be moved by female tract mechanisms, but we rarely understand which sex is controlling sperm dispersal, and how, where and when this occurs through the whole process. Several female mechanisms could control sperm transfer, progress and activity; from mechanical contractions and hydrostatic pressures in the female tract, to sorting sperm from different males in designated organs, and through completely ejecting ejaculates or exerting spermicidal actions (Firman et al., 2017). Biochemical complexity in which these dynamics take place is also important, with evidence that the female tract can be either supportive or, at times, hostile to certain male gametes (Firman et al., 2017; Wolfner, 2011). Ostensibly, much remains to be discovered about this reproductive diversity, with recent *in vivo* research using GFP-tagged sperm revealing high levels of activity and interaction between sperm from different males and different areas of the female tract (Manier, Belote, et al., 2013; Manier, Lüpold, et al., 2013).

External fertilization, in which gametes fuse outside the body in an aqueous environment, appears to present far fewer opportunities for females to exert post-mating control over fertilisation. Interactions between gametes cannot benefit from a complex reproductive tract with opportunities for differential sperm uptake, storage, and management. However, despite its increased reproductive ‘simplicity’, studies have shown that external fertilization can indeed allow cryptic female choice via adaptations that encourage the ‘right’ sperm, or discourage the ‘wrong’ sperm, to fertilize (Firman et al., 2017). For example, gamete recognition systems in or on the egg, and reproductive fluids that are released with the eggs, are known to influence sperm behaviour and fertilisation outcome (Evans et al., 2013; Yeates et al., 2013). It is the relative simplicity of these systems compared to internal fertilizers, and the tractability of external fertilisation for controlled *in vitro* fertilization experiments, that have enabled significant advances in understanding the outcomes and potential mechanisms that control sperm-egg interactions in the context of post-mating selection from sperm competition and cryptic female choice.

Some of our most fundamental knowledge about sperm-egg interactions comes from broadcast-spawning marine invertebrates. The associations between bindin molecules (Palumbi, 1999), and between lysin and its vitelline envelope receptor (VERL) (Swanson & Vacquier, 1997), have been described in detail in sea urchin and *Haliotis* respectively, where biochemical mechanisms control against the risk of heterospecific sperm attachment or egg membrane penetration (Metz et al., 2016; Palumbi, 1999), influencing individual fertilisation success (Hussain et al., 2016). Similarly, more recent work has described the mechanisms by which female-derived chemoattractants within egg-associated reproductive fluids mediate post-mating mate choice, fertilization success and offspring fitness in mussels (Fitzpatrick et al., 2012; Oliver & Evans, 2014). In fish, females manufacture ovarian fluid, which is released into the coelomic cavity with maturing eggs (Hirano et al., 1978). It contains a complexity of nutrients, metabolites and hormones (Hirano et al., 1978; Ingermann et al., 2001; Lahnsteiner et al., n.d.), and once spawned shows the highest concentration in proximity to the micropyle entrance of eggs. Ovarian fluid identity of different females has been found to differentially impact sperm swimming behaviour (Alonzo et al., 2016), and influence fertilisation outcome according to the genetic relatedness of males (Butts et al., 2012; Gasparini & Pilastro, 2011) and their spawning origin (Beirão et al., 2014). In salmonids, ovarian fluid comprises up to 30% of the spawned egg mass, and its influence on sperm is relatively well studied (Galvano et al., 2013; Johnson et al., 2020; Purchase & Rooke, 2020; Turner & Montgomerie, 2002; Zadmajid et al., 2019). There is increasing evidence that this reproductive fluid can act as a ‘fertilisation filter’ for or against sperm from different partners, enabling cryptic female choice. This facilitates sperm selection even in highly polyandrous externally fertilisers like Atlantic salmon (*Salmo salar*), where a single egg batch can be sired by up to 16 fathers (Weir et al., 2010). Yeates et al. (Yeates et al., 2013) showed that ovarian fluid allowed females to apply conspecific sperm precedence when facing *in vitro* hybridization risks between Atlantic salmon and brown trout (*Salmo trutta)*. However, we do not yet know the exact mechanisms facilitating such choice.

Sperm swimming propulsion is created by the flagellum, whose function is influenced by chemical (J. Cosson, 2015; Kholodnyy et al., 2019) and physical (J. Cosson, 2015; J. Cosson & Prokopchuk, n.d.; Holwill, 1977) conditions. The different responses of sperm behaviour reported in presence of ovarian fluid, and their resulting effects on fertilization (Alonzo et al., 2016; Galvano et al., 2013; Gasparini et al., 2012; Rosengrave et al., 2009), have been associated to changes in pH (Wojtczak et al., 2007), ionic composition (Rosengrave et al., 2009), and viscosity (Turner & Montgomerie, 2002) that control flagellar beating (Kholodnyy et al., 2019). While the effects of chemistry (Rosengrave et al., 2009; Wojtczak et al., 2007) and temperature (Dadras et al., 2016, 2017) have been more frequently investigated (J. Cosson, 2015; Dadras et al., 2016; Kholodnyy et al., 2019), the influence of changes in viscosity on swimming sperm remain poorly explored in external fertilizers (Kholodnyy et al., 2019; Lauga, 2007). There is evidence that fish ovarian fluid possesses structural properties that makes for a non-Newtonian viscous response (where viscosity changes depending on the force applied) that is very different to water (Rosengrave et al., 2009), and this peculiar viscous response could influence the biophysics of sperm swimming behaviour in external fertilisation environments. To describe such function, we conducted detailed measurements of its biophysical characteristics using a rheological approach commonly used in soft-matter physics. We sought to uncover the rheological nature of ovarian fluid when different forces are applied to it, thus exploring how its non-Newtonian behaviour could affect sperm activity, penetration, bioenergetics, and guidance to fertilisation in a context of sperm competition and cryptic female choice.

## MATERIAL AND METHODS

### Sample collection and preliminary measurements

Wild anadromous Atlantic salmon were collected in early September from a fish ladder at Grand Falls (48° 55’ N, −55° 39’ W) during their up-stream spawning migration on the Exploits River (Newfoundland, Canada). Following previous protocols (Rooke et al., 2019), fish were transferred to covered, outdoor tanks next to the river, and experienced ambient temperatures and light. Over two weeks in early November, females were assessed for ovulation using gentle abdominal pressure, fish were then anaesthetised using a solution of 2ml/L clove oil, measured for length and weight, and stripped of eggs after drying the urogenital pore. Each female’s eggs (and associated ovarian fluid) were kept in sealed glass jars, enclosed with bubble wrap, and placed in a cooler of wet ice for transport to the laboratory. Each egg batch was separated from its ovarian fluid using a fine mesh net (Purchase & Rooke, 2020) within 10 hours of stripping. For each ovarian fluid we recorded volume and weight to deduce density, followed by pH and conductivity.

### Rheological characterization of ovarian fluid

The mechanical properties of many soft biological materials are neither purely viscous (liquid-like) nor purely elastic (solid-like), and these rheological properties correlate strongly with their function (Storm et al., 2005). Structured fluids often do not flow until they reach a critical stress level, below which a material is considerable elastic, and above which the structure of the material breaks down and starts to flow. Two experiments were performed to define how the ovarian fluid’s polymeric structure (and related physical properties that in turn would affect sperm swimming activity) can be modulated, depending on swimming sperm flagellar beat frequency. Specifically, we tested ovarian fluid ‘behaviour’, both under steady shear (i.e., “flow curves”) and under small-amplitude oscillatory shear (SAOS). The former examines the viscoelastic response of the ovarian fluid by continuous deformation and breakup of internal networks, while the latter can probe weaker internal structures (Bird et al., 2021; Markovitz, 1981). A preliminary rheological analysis (n= 5 fish) was conducted to assess different fluid preservation methods (see supplementary material). Each frozen sample was thawed at room temperature for 1hr prior to analysis, and measurements were made using 1.5 mL aliquots. All the analyses were performed in the Soft Matter Lab at Memorial University using an MCR 301 rheometer, equipped with a cone-plate (CP50-0.5, 50 mm diameter plate and cone angle, Anton Paar GmbH, St. Albans, UK) system. Ovarian fluid samples were individually filtered through a 200 μm sieve to remove any particulates (e.g., coagulated blood, ovarian tissue) that could influence the rheological measurements. Pipetted fluid was equilibrated for three minutes at the plate temperature of 6°C, allowing for homogenous sample relaxation from any uncontrolled pre-shear imposed on the fluid during loading.

### Steady-state shear properties

Samples were tested for their resistance to flow in order to measure their viscosity under a specific rate of deformation. To obtain a flow curve, the shear stress was measured for a range of shear rates 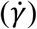, from 10 to 500 s^-1^ in 50 equally spaced steps. The resultant shear stresses of the ovarian fluid were measured to determine the *apparent* viscosity η_a_, which was averaged across three aliquots per female (n= 11) and plotted as a function of the shear rate.

Among each of the three ovarian fluid aliquots per fish, a run with distilled water was performed as a control. For distilled water (pure Newtonian fluid), a theoretical positive relationship between shear stress and shear rate should be linear and the fit line should pass through zero. When the profiles of water runs were fitted, a positive intercept (typical for these kind of measurements) of 0.0133 Pa was concluded to be low-shear rate instrumental noise. It subtracted from all the water and ovarian fluid samples as standardization (shear stress −0.0133 Pa)/(shear rate), creating a small change in values. A comparison of individual ovarian fluid viscosity profiles with distilled water for each of the instrumental replicates allowed us to assess variability among females.

The apparent viscosity of ovarian fluid decreased with increasing shear rates, in contrast with water whose apparent viscosity (η_a_= 0.00151 ± 0.00003 Pa·s) was independent of shear rate. The apparent viscosity at 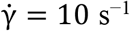 was roughly 10 times the viscosity of water but returned comparable at 100 s^−1^ (see Results). For three females the ovarian fluid samples had apparent viscosities η_a_ in the order of 0.003 Pa·s at 10 s^−1^, showing no meaningful differences with the rheological behaviour of water at the same shear rate. Likely, these samples were contaminated by urine and/or water during stripping of gametes and for these reasons were not included in the main results. The remaining 11 flow curves were globally fitted to the form 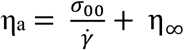, which is a simple equation incorporating an elastic component, the yield stress *σ*_00_ which must be overcome before there is flow, and a viscous component η∞, which represents the viscosity at very high shear rates. This simple form was arrived at when fits to a more complicated formula, the Herschel-Bulkley equation 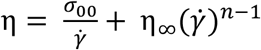 (Herschel, 1926) resulted in power laws *n* that were very close to unity.

### Small Amplitude Oscillatory Sweeps

To preserve finer polymeric structures and obtain a dynamic profile that informs about the viscous and elastic components, we subjected the ovarian fluid to small-amplitude oscillatory shear. For these measurements, a sinusoidal deformation (γ = γ_0_ sin ωt) was imposed on the sample at a fixed frequency ω and a maximum amplitude (γ_0_) (Schoff & Kamarchik, 2005). Measurements were performed for a range of frequencies (ω), from 0.01 to 500 rad· s^−1^ in 24 equally spaced logarithmic increments. The storage modulus,

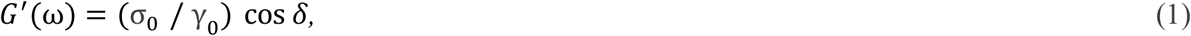

and the loss modulus,

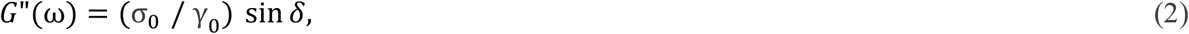

were obtained as a function of frequency (ω). The modulus of the complex viscosity η* was obtained from the relation

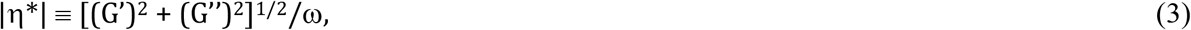

while the damping factor (or loss factor) tan δ ≡ G”/ G’ represents the ratio between viscous and elastic contributions to the viscoelasticity.

### Applicability of the Cox-Merz rule

The Cox-Merz rule, an empirical method to rationalize steady shear and oscillatory rheological data (Cox & Merz, 1958), was used to compare the two different rheological analyses adopted in our study. A strong correlation between two independent methodologies is a good consistency check. This rule states that the apparent viscosity 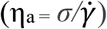 at a specific shear rate 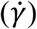 is equal to the complex viscosity (|η*(ω)| = |*G*^*^(*ω*)|/*ω*) at a specific oscillatory frequency (ω), that is

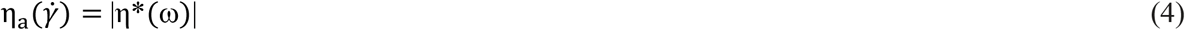

When the rule is obeyed, rheological properties of a fluid can be described by either oscillatory or steady-state shear experiments (“Engineering Properties of Foods,” 2014).

### Statistical analyses

All ovarian fluid measurements and fish morphological data (mean ± SD, 95% CI and Coefficient of Variation (CV%)) were summarised using the descriptive statistics function in GraphPad Prism, version 8.0.0, (GraphPad Software, San Diego, California USA). Rheometer reads were first standardized for instrumental error and the model fits were applied as described above. Subsequently, the average values of *G’* and *G”* (dependent variables) across all the sampled females were pair-wise compared trough t-tests at specific frequencies (independent variables) of interest within two shear stress ranges, 0.001 to 0.105 and 0.105 to 1 rad· s^−1^, to double-check their uniformity within the plateau region and/or alternatively the prevalence of either the viscous or the elastic component of the ovarian fluid in this dimensional range. Normality of the residuals was ensured through D’Agostino-Pearson test followed by Shapiro-Wilk test (P= 0.2174 and 0.4697, respectively). Throughout the analyses, the statistical significance threshold used was α= 0.05.

## RESULTS

Ovarian fluid characteristics varied among individual females (Table 1). For context, coefficient of variation ((standard deviation / mean)*100) of fish length was 10% while body weight (which included eggs and ovarian fluid) was 34%. The amount of ovarian fluid produced for a given size of fish or mass of eggs was very inconsistent among females (CV ∼50%). Conversely, fluid density, pH and conductivity were similar (<10%, and thus less variable than fish length). Apparent viscosity was highly variable among fish, but all exhibited clear non-Newtonian behaviour. The amount of variation declined with the shear rate applied, being CV=57% among females measured at 10 s^-1^ and CV=17% at 500 s^-1^ (Table 1, Figure 1).

**Table 1.** Rows group wild Atlantic salmon female growth-related parameters (length (cm) and weight (Kg), ovarian fluid (OF) pH, conductivity (mS cm^-1^), density (g/cm^3^), volume (mL; per centimetre, per kilogram and per 10 grams of eggs) and apparent viscosity values at 10, 50, 100 and 500 rad• s^-1^ (Pa• s). In columns, left to right values are expressed as means ± standard deviations (SD), range (min-max), and coefficient of variation (CV %) among females (n=11).

| | Mean<br>± SD | Range<br>(Min-Max) | CV<br>(%) |
| --- | --- | --- | --- |
| Fish length (cm) | 54.55 ± 5.714 | 40.90 - 74.20 | 10% |
| Fish weight (Kg) | 1.556 ± 0.530 | 0.57 - 3.54 | 34% |
| OF volume (ml) per cm of fish | 1.06 ± 0.54 | 0.23 - 2.20 | 51% |
| OF volume (ml) per Kg of fish | 37.78 ± 19.05 | 7.97 - 64.98 | 50% |
| OF volume (ml) per 10 gr of eggs | 2.38 ± 1.32 | 0.53 – 4.41 | 55% |
| OF pH | 8.264 ± 0.117 | 8.010 - 8.57 | 1% |
| OF conductivity ($\text{mS} \cdot \text{cm}^{-1}$) | 14.19 ± 0.674 | 11.77 - 15.19 | 5% |
| OF density ($\text{g} \cdot \text{cm}^{-3}$) | 0.993 ± 0.091 | 0.8220- 1.530 | 9% |
| OF apparent viscosity ($\text{Pa} \cdot \text{s}$) at $10\text{ rad} \cdot \text{s}^{-1}$ | 0.012 ± 0.006 | 0.006 - 0.029 | 57% |
| OF apparent viscosity ($\text{Pa} \cdot \text{s}$) at $50\text{ rad} \cdot \text{s}^{-1}$ | 0.004 ± 0.001 | 0.002 - 0.008 | 40% |
| OF apparent viscosity ($\text{Pa} \cdot \text{s}$) at $100\text{ rad} \cdot \text{s}^{-1}$ | 0.003 ± 0.001 | 0.002 - 0.005 | 32% |
| OF apparent viscosity ($\text{Pa} \cdot \text{s}$) at $500\text{ rad} \cdot \text{s}^{-1}$ | 0.002 ± 0.000 | 0.001 - 0.003 | 17% |

**Figure 1.**
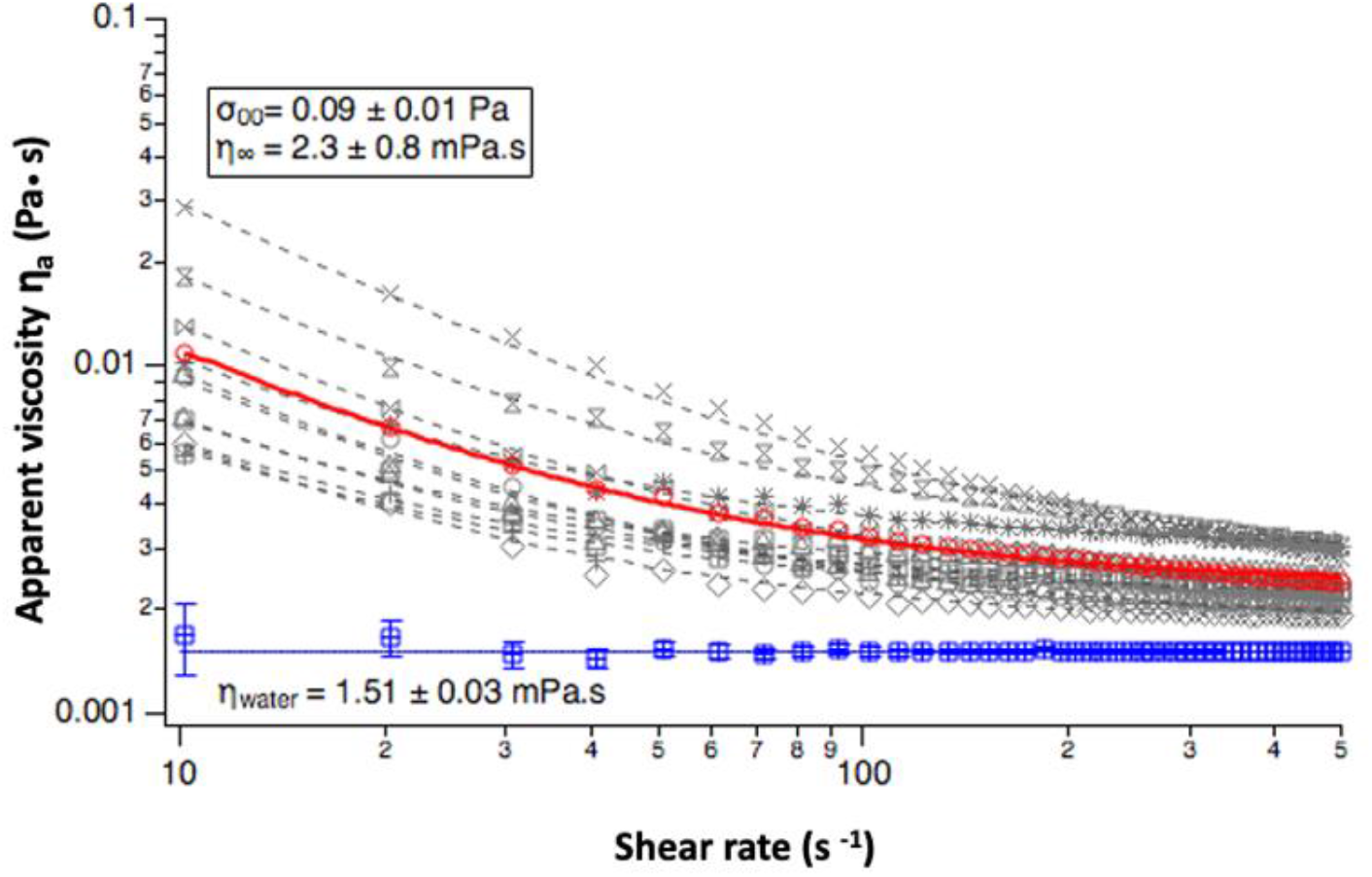
Apparent viscosity obtained from the steady shear flow curves (η) of Atlantic salmon ovarian fluid samples (n= 11, in grey) and water controls in blue, plotted versus shear rate (s^-1^) on a log-log scale. Grey symbols and dotted lines represent individual ovarian fluid means across 3 replicates per female and their fitted equations respectively, while the red and blue symbols and the continuous lines represent the mean across all ovarian fluid samples (red) and water controls (blue). The symbols σ_00_ and η_∞_ are respectively the Yield stress and the apparent viscosity at high shear rates obtained from the fitting to the Herschel-Bulkley equation; η_water_ instead represents the average apparent viscosity value of water within the analysed shear rates (mean ± SD).

### Ovarian fluid rheology in steady-state shear flows

To measure the viscosity under a linearly increasing rate of deformation, the ovarian fluid samples were tested for their resistance to flow for a range of shear rates (10 to 500 s^-1^). The resulting shear stress responses from the deformed ovarian fluid were measured to determine the apparent viscosity of the material at each of the measuring points.

Atlantic salmon ovarian fluid showed non-Newtonian shear thinning behavior indicating successive loss of polymer entanglements with increasing shear rates (Figure 1). The Herschel-Bulkley equation fits (Figure 1) returned a mean value of yield stress *σ*_00_= 0.09 (± 0.01 Pa) and a mean value of the high-shear viscosity η_∞_= 2.3 (± 0.8 mPa • s) with the ovarian fluid showing an average 97% decline in viscosity as an increasing shear rate was applied through the rheometer’s plate. Variability among all females and water control are also shown in Figure 1.

### Small Amplitude Oscillatory Sweeps and dynamic shear properties of the ovarian fluid

The dynamic viscoelastic behaviour of the ovarian fluid dispersions was also determined by applying small amplitude oscillatory shear (SAOS) frequency sweep. The storage modulus *G*’ and loss modulus *G’’*, shown in Figure 2(A), were not different at low frequencies, with both having a value of approximately 0.1 Pa in the 5 measuring steps between 0.01 and 0.105 rad· s^−1^ (0.065 ± 0.011 Pa and 0.077 ± 0.001 Pa respectively (mean ± SD); P≥ 0.05, t= 2.77, df=4) and describe a pure viscoelastic fluid where the elastic and the viscous components of the fluid are comparable. Both *G’* and *G’’* decreased slightly between 0.01 and 0.07 rad· s^−1^ (note the Log_10_-Log_10_ axes) and thereafter maintained constant plateau values until the shear rate reached 1 rad s^−1^. Note that this plateau value is numerically proximate, given the errors, to the value obtained for the yield stress in the steady shear measurements. Salmon ovarian fluid is therefore a gel-like structure at low frequencies and becomes more dominantly liquid-like at frequencies higher than 10 Hz. Interestingly, this structural shift occurs in a dimensional range that overlaps with the frequencies exerted by salmon sperm when swimming through the ovarian fluid to reach the egg (refer to dashed vertical lines in Figure 2A, B). This is confirmed also by the fact that at low frequencies, the gel-like structure is supported by a value of tan δ = *G’’/ G’* of 1 (crossover or gel point, See Fig 2B), however between frequencies from 0.10 to 1 rad· s^−1^ (6 steps) the loss modulus *G’’* (mean 0.081 ± 0.02) was marginally higher (P<0.001, t= 32.93, df= 5), than *G’* (0.052 ± 0.01). As observed through the study of their first and second derivatives, *G’* and *G’’* trends start to slowly diverge, more intensely from 10 Pa (at 1.59 Hz) onward revealing a breakpoint in the polymer that exacerbates together with increasing shearing rates (Fig 1, 2 in supplementary material). Specifically, the storage modulus reached 0 Pa between 47.6 and 312 rad· s^−1^ (7.58 and 49.66 Hz), showing that the elastic response of the polymer under these frequencies is null (liquid-like); and viscous forces at their maximum in this frequency range instead prevailed. As a result, the absolute value of the complex viscosity (|η*|) decayed until reaching its a minimum of 0.005 Pa·s, at a frequency near 8 Hz (50 rad· s^−1^), (see Fig 2A). Interestingly, |η*| increased after this measuring point. Values of tan δ = *G’’/ G*’, were similar at low frequencies and also showed a clear dependence in the same frequency range increasing to 34 ± 17 at the highest frequencies.

**Figure 2.**
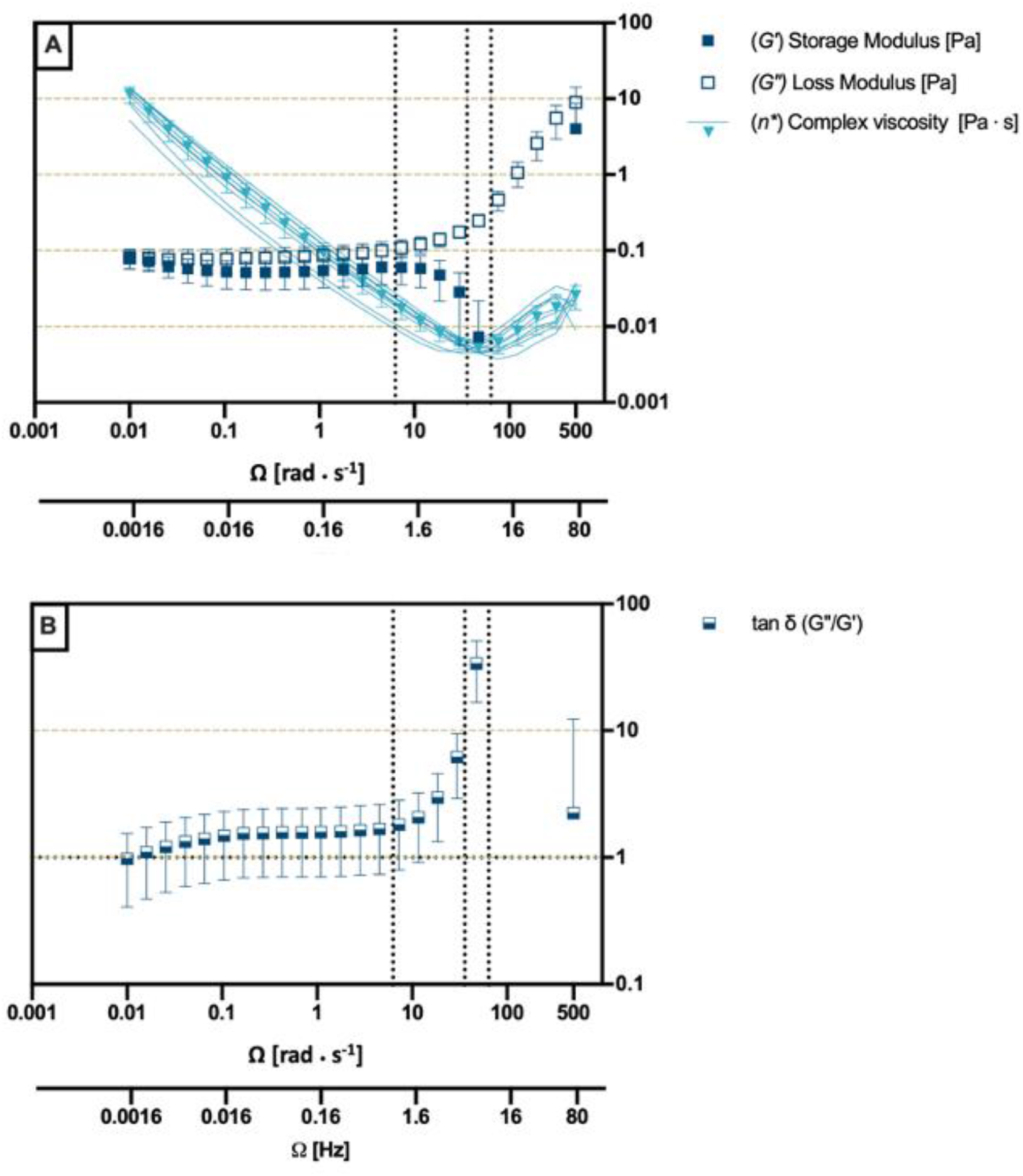
**(A**) Storage modulus (G’), loss modulus (G’’) and complex viscosity (η*) of Atlantic salmon ovarian fluids (n= 11), to describe the relation between the viscous and elastic components of the fluid at increasing angular frequencies (0 <Ω< 500 rad· s^−1^). Data are presented as means ± SD, continuous lines for η* represent individual ovarian fluid means across three replicates for each female). **B)** Loss factor (tan δ= G”/G’) of Atlantic salmon ovarian fluids (mean ± SD) plotted versus frequency (rad· s^−1^, Hz for reference), where tan δ= 100 for a liquid material with a pure viscous behaviour and tan δ= 0.01 for a solid material with an ideally elastic behaviour. Vertical dotted lines from left to right represent a reference baseline at 1 Hz and Atlantic salmon average sperm tail beat frequencies in Hz from Dzievulska et al., 2011a, b. Graph is plotted on log-log scale.

### Comparison of steady and oscillatory shear

The steady state properties of the ovarian fluid were compared with the dynamic states by applying the Cox-Merz rule. This rule, applied to polymers, enables the identification of secondary flow behaviours and/or breaking down of the fluid’s polymeric network under a certain imposed stress. Apparent viscosities (η_a_) obtained in the flow curves, and absolute values of complex viscosities (|*η*\*|) resulting from the small amplitude oscillatory sweep experiments, were plotted as a function of shear rate rad s^−1^, fitted to the best trend and assessed for deviations between the curves’ profiles (Figure 3). Ovarian fluid η and *η*\* followed the same trend with many remarkable similarities. When the oscillatory shear probed lower frequencies, the curves overlapped very closely between 10 and 50 rad· s^-1^. From the steady shear results, we extracted a yield stress *σ*_00_ = 0.09 ± 0.01 Pa, which is close to the *G’* plateau value of *σ*_0_ = 0.068 ± 0.006 Pa. However, above 50 rad· s^−1^ (8 Hz), there is an increase in |*η*\*|.

**Figure 3.**
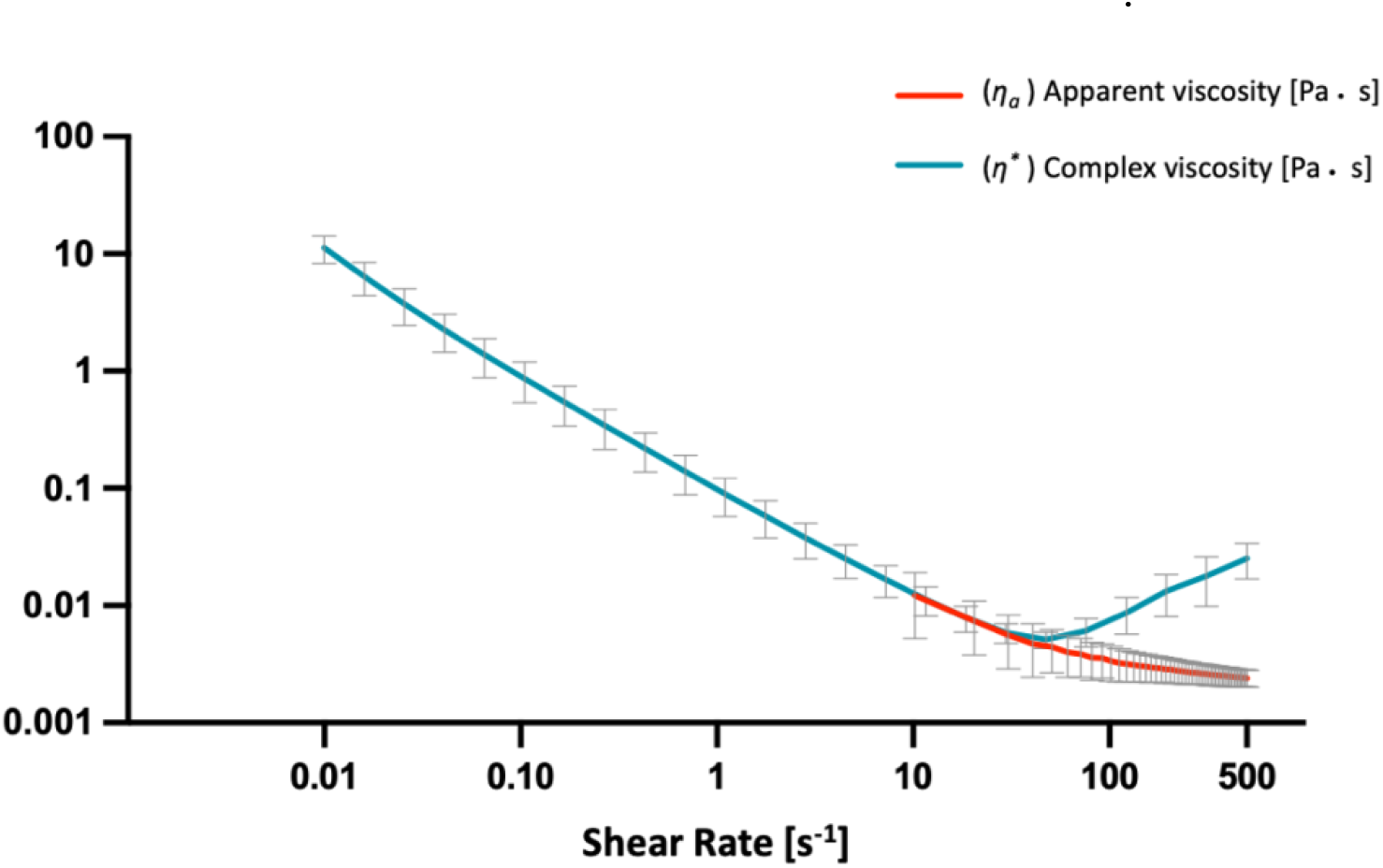
Comparison between the steady state and the dynamic properties of Atlantic salmon ovarian fluid (OF). Apparent viscosity obtained from the steady shear flow curves (η_a_) of Atlantic salmon ovarian fluid (OF) samples (N= 11, red) and complex viscosity (η*) of Atlantic salmon ovarian fluids (n= 11, blue) plotted versus shear rate (s^−1^). Values are presented as mean ± SD (grey vertical bars). Graph is plotted on log-log axes.

Beyond this frequency, the Cox-Merz rule was not obeyed, meaning that η_a_ and |η*| values obtained at a specific shear-rate are not equal when compared between the two different methodologies used. It should be noted that steady shear is much more disruptive to the gel structure than oscillatory shear. Thus, while we must be cautious with interpreting the rise in |η*| between 50 and 500 s^−1^, it is nevertheless feasible that this rise is indicative of a rise in the SAOS viscosity.

## Discussion

We describe the rheological characteristics of Atlantic salmon ovarian fluid to understand the possible involvement in sexual selection mechanisms. We subjected ovarian fluid from different females to both variable shear stresses (steady-state rheology) and angular frequencies (small-amplitude oscillatory sweeps), similar to those exerted by sperm swimming through ovarian fluid to fertilize eggs (Dziewulska, Rzemieniecki, & Domagała, 2011; Dziewulska, Rzemieniecki, Czerniawski, et al., 2011). This allowed for the identification of the main viscoelastic profile of the fluid, but also for inferring secondary flow behaviours and the eventual breaking down of macromolecular entanglements under a certain imposed stress. In particular, small-amplitude oscillatory measurements (SAOS) describe the viscous and elastic components within the ovarian fluid that could affect fertilisation dynamics. We found that the physical characteristics of salmon ovarian fluid clearly show a non-Newtonian viscoelastic nature, where shear-influenced changes in viscosity and elasticity might have the potential to influence fertilization. Here, we discuss the structural characteristics of the ovarian fluid that could influence sperm and explore the potential of its non-Newtonian properties to be adaptive.

### Shear-thinning behaviour in steady-state rheology and under small amplitude oscillatory sweeps

Our results indicate that ovarian fluid, which is a gel at its relaxed state (between solid- and liquid-like behaviour), is a shear thinning viscoelastic-liquid at low frequencies, and may exhibit a shear thickening phase at high frequencies. This shift from gel to a more passable medium, together with minimum viscosity values observed within the range of average beating frequencies exerted by salmon sperm, points out an interesting overlap that might be linked to ‘bio-mechanical co-evolution’ of female and male gametes. Specifically, Atlantic salmon ovarian fluid has a viscosity at its relaxed state that is on average 60 times that of water, being 0.09 Pa. A hypothetical beating frequency of 1Hz would yield to the absolute value of the complex viscosity |*η*\*| of 0.017 Pa·s (five times lower than at its relaxed state), while a beating frequency of 12 Hz yields 0.006 Pa·s. Sperm movement occurs at low shearing rate (Brokaw, 1965, 1966) and the reported sperm beat cross frequency (BCF) values present in literature (∼ 5-10 Hz) (Dziewulska, Rzemieniecki, & Domagała, 2011; Dziewulska, Rzemieniecki, Czerniawski, et al., 2011) are in a similar range of frequencies as used in our experiment. Fascinatingly, these frequencies correspond to either the shear thinning region or to the minimum values of apparent and complex viscosity reported, having a meaningful biological translation. Another intriguing possibility is that the departure from the Cox-Merz rule at high frequencies might actually signal an increase in viscosity when measured in a sensitive way (SAOS) that does not disrupt gel structure, but not when measured in a more disruptive manner (steady shear). There might therefore even exist an optimal beat frequency window below *and* above which the ovarian fluid is effectively more viscous.

Flagellar beating frequency varies considerably with temperature, pH, time, activation medium (e.g. water vs. ovarian fluid) and methodology used to detect it (Cosson, 2021; Zadmajid et al., 2019). Measures from other salmonids obtained at higher temperatures, in a diluted solution of ovarian fluid, and using stroboscopic techniques, show higher frequencies, such as ∼50 Hz for *O. mykiss* (M. P. Cosson et al., 1985) and ∼80 Hz for *O. tshawytscha* (Butts et al., 2017). For this reason, the authors will more cautiously consider for this discussion a broader sperm beating frequency of salmon sperm in ovarian fluid going as up as 80 Hz.

Our viscosity measures are considerably higher than reported for other fish species, such as 0.0038 Pa.s (2.76 times that of water) for Arctic charr (*Salvelinus alpinus*) when measured at 0.5 Hz under a plate viscosimeter (Turner & Montgomerie, 2002). In chinook salmon (*O. Tshawytscha*) ovarian fluid viscosity decreased from 0.0042 to 0.0027 Pa·s as shear was increased from 7 to 72 Hz (Rosengrave et al., 2009). These lower values reported in other species might be related to the higher starting frequency used as compared to ours. In fact, if paralleled to what we found in *S. salar*, the starting point of 7 Hz used for *O. tshawytscha* falls within the shear thinning phase of the fluid, implying that this was first probed already under a certain initial stress rather than at its relaxed state, thus masking a potentially higher relaxed state viscosity. In our case, by controlling for instrumental uncertainty and comparing two different rheological approaches, we had the advantage to precisely probe and extract realistic zero shear viscosities and low shear values. This is relevant because our results evidence not only that the gap in viscosity caused by increased shear is greater than previously thought, but so will be the biological implications resulting from different frequencies shearing the fluid.

### Viscous and elastic components within the ovarian fluid that could affect fertilization dynamics

Viscous compounds were already known to influence the flagellum, resulting in a lower velocity (Brokaw, 1965). Brokaw (Brokaw, 1966, 1983) investigated sperm flagellar behaviour in response to increased viscosity in three marine Phyla (Anellida, Tunicata and Echinodermata), finding a decrease of both beat frequency and wavelength, similar to what was found in chinook salmon (Butts et al., 2017). These authors partially justified an observed increase in velocity and propulsive efficiency of sperm swimming in ovarian fluid through the non-Newtonian properties of this medium. These were firstly described in a study by Rosengrave and colleagues (Rosengrave et al., 2009), who explored its response to shear rates under a constant rotational force (steady state properties). By including both steady state measurements and SAOS, we add crucial information on the specific elastic and viscous components within the ovarian fluid that could justify the changes in sperm behaviour reported by other authors and further investigate its role during reproduction.

In view of our rheological results which show viscoelastic behaviour of salmon ovarian fluid, new considerations need to be made because the viscous (liquid-like) and elastic (solid-like) components of the fluid defining its changing complex viscosity cannot be neglected when analysing sperm energetics and outcome. In studies with internally fertilizers (mammals), viscoelastic reproductive fluids have been found to decrease spermatozoa velocities as viscosity increased while concurrently increasing their linearity (Suarez & Pacey, 2006). *Bos taurus* have higher thrust efficiencies of sperm when swimming in a non-Newtonian fluid rather than in a Newtonian one, which has been suggested might be due to a better energetic exploitation of the elastic responses of the fluid (Hyakutake et al., 2019). In our case, we observe a drop in absolute value of the complex viscosity as the frequency is increased up to 8 Hz (absolute viscosity minimum) when subjecting salmon ovarian fluids to SAOS, suggesting that until this point sperm find an increasingly thinner polymeric network that gets looser with frequency. This happens first in presence of a good elastic component that instead collapses in the ‘armpit region’, having the potential to positively influence sperm linearity and guidance. In this fluid, sperm with different tail beating frequencies, would in principle face substantially different polymeric structures within the shear-thinning phase. This shear-thinning flow behaviour could either facilitate sperm getting into the egg, or it could also enable cryptic female choice if a specific sperm, its morphological phenotype, swimming behaviour or another other trait, is favoured over the one of a rival male competing to fertilize the eggs. Moreover, if considering the reported within-male sperm variability observed in *S. salar* (Immler et al., 2014), it is presumable that the physical properties of ovarian fluid might have a role also in within-male sperm selection.

Sperm traits are under strong selection (Fitzpatrick et al., 2020; Fitzpatrick & Lüpold, 2014), with recent studies evidencing a relation between some sperm traits and offspring fitness (Immler et al., 2014) and a correlation between sperm phenotype and genotype (Alavioon et al., 2017). Moreover, sperm within the same ejaculate can experience different stressors that negatively affect their swimming behaviour; the impairment of these ‘abnormal’ gametes is also reflected on a molecular level (e.g., DNA fragmentation) (Fernández et al., 2003), influencing the quality of the information transmitted to the zygote and accordingly its performance. Flagellar activity declines with time post-activation, while the osmotic- and ROS-derived damage experienced by the sperm cell increases (Kholodnyy et al., 2019). Therefore, the peculiar non-Newtonian properties of this fluid, shear-thinning at low shear rates, followed possibly by shear-thickening, might help select the best performing sperm also within a single ejaculate with the objective to limit the chance of ‘abnormal’ sperm getting into the eggs. The frequency-dependent minimum in viscosity, raises therefore the intriguing possibility that the ovarian fluid selects for an optimal speed, providing a viscosity cost for both slow and fast beating sperm. Augmenting the swimming cost also for an ultra-fast fertilization could eventually allow selection based on further biochemical mechanisms, that are known to be pivotal for sperm egg interaction, can influence the reproductive outcome and have been advocated in reducing the hybridization risk with other species (Yanagimachi et al., 2017).

It is well accepted that the guidance within reproductive fluids occurs by means of chemical and biochemical cues that can differentially enhance the reproductive outcome from different males as demonstrated in a range of external fertilizers (Cosson, 2015; J. Cosson et al., 2008; Evans & Lymbery, 2020; Evans & Sherman, 2013; Kholodnyy et al., 2019; Yanagimachi et al., 1992; Zadmajid et al., 2019). We propose that more consideration should be given to the physical characteristics of the ovarian fluid that could affect sexual selection processes. Females might be able to facilitate the progression of the high quality and fast beating sperm, within and among ejaculates. Also, in view the variability observed across females; it could be suggested that these might have different capabilities to exert this selective potential, and is not to exclude how such potential could change with the hydration grade of the ovarian fluid as the reproductive season advances.

### Shear-thickening behaviour

Under SAOS at the highest shear rates measured we observed a significant stiffening of the polymer. This did not occur in steady state measurements, where the fluid continued to thin up to 80 Hz. This difference could identify the presence of weak network associations that are broken in steady state flow measurements, where a continuous rotational force is applied on the fluid. In contrast, these interconnected networks are unaffected in the oscillatory shear tests. In this case, the ovarian fluids were subjected to sinusoidal shear stresses within a linear range small enough that the macromolecular entanglements are preserved. A shear thickening phase at very high frequencies, mostly in absence of any elastic component, would suggest that sperm swimming efficiency could be exclusively dependent on its speed and on the fluid viscosity, without exploiting the positive effects on linearity that some elasticity would provide. The lack of elasticity on the other hand may also be promoting a more circular swimming pattern, which in a closely related species *(Oncorhynchus mykiss)* has been linked to augmenting the chances of fertilizing the egg (Wojtczak et al., 2007).

It could be further speculated that a shear thickening phase at high frequencies might also be linked to the ‘necessity’ for ovarian fluid to stay close to the eggs and not be washed away – an aspect of natural selection. Salmon spawn in rivers and an infinitely shear-thinning ovarian fluid would enhance its chances of being dispersed and diluted very quickly, thus depriving the eggs from its known beneficial effects on fertilization (Alonzo et al., 2016; Butts et al., 2012; Gasparini & Pilastro, 2011; Poli et al., 2019; Yeates et al., 2013). Other works have shown that a shear-thickening behaviour observed at high frequencies could be also derived from inertial forces. For example, a study of hagfish slime (Böni et al., 2016) presented similar behavior at low frequency with G”/G’ of order one, indicating an ultra-soft material having weak elastic properties. However, in that study a rise in G’’ (and drop in G’) at higher frequencies was attributed to instrument inertia. We cannot exclude that this could also be the case here. Moreover, although the idea that the ovarian fluid may have a natural selective function at very high shear rates is indeed fascinating, this specific aspect lies outside the scope of this study and was not tested specifically. Future experiments should try to provide insights in this regard by testing the ovarian fluid dispersion capacity from eggs under different shear rates that could better simulate the riverbed waterflow. Additionally, it could be also tested whether the shear-thickening phase observed at the upper end of our analysed range is sincere and if this persists at very high frequencies with a beneficial effect on the eggs (e.g., higher diffusion, mechanical resistance, pathogen barrier) (Elofsson et al., 2003).

### Conclusive remarks and future perspectives

Ovarian fluid physical properties deserve more attention and considerations when studying processes of sexual selection such as selection on sperm performance, sperm competition assays and fertilization trials, both *in vivo* and *in vitro*. The characteristic rheological behaviour of the ovarian fluid we report here underlines the importance of including it as a preferred sperm activation medium over pure water to simulate a more natural fertilisation environment and benefit from its effects on sperm.

Our discovery yields a number of predictions to be tested in the future, including testing whether the physical properties of ovarian fluid act as a filter for specific sperm or, whether its structure only ameliorates sperm performance in general. Notably, our findings suggest that processes enabled by non-Newtonian reproductive fluids within female internal genital tracts, like lubrication, facilitation and capacitation, should also be applied to the external fertilization environment. This opens new avenues into the study of cryptic female choice with important implications for understanding the evolution of sexual traits and exploring the underestimated role of physical properties of the fertilization environment that surrounds the gametes both in nature and in artificial fertilization protocols.

## ACKNOWLEDGEMENTS

We would like to dedicate this paper to Matthew Gage, who prematurely passed away during the final drafting of the manuscript, but whose valuable contribution heavily influenced its framework. This research was supported by grants to Craig Purchase from the Natural Sciences and Engineering Research Council of Canada, the Canada Foundation for Innovation, the Research and Development Corporation of Newfoundland and Labrador, and the Atlantic Salmon Conservation Foundation. Thanks to the staff at the Environmental Resources Management Association for acquiring the fish for us, and to Margaret Litt, Steven Poulos and Ryan Carrow for assistance with sampling. We express gratitude toward Tatzuo Izawa from the Soft Matter Lab for the technical support provided and for the insights given during preliminary phases of this study.

## Notes

### Competing Interest Statement

The authors have declared no competing interest.

### Summary of Updates

Supplementary files were added

## References

Alavioon, G., Hotzy, C., Nakhro, K., Rudolf, S., Scofield, D. G., Zajitschek, S., Maklakov, A. A., & Immler, S. (2017). Haploid selection within a single ejaculate increases offspring fitness. Proceedings of the National Academy of Sciences of the United States of America, 114(30), 8053– 8058. https://doi.org/10.1073/PNAS.1705601114/-/DCSUPPLEMENTAL

Alonzo, S. H., Stiver, K. A., & Marsh-Rollo, S. E. (2016). Ovarian fluid allows directional cryptic female choice despite external fertilization. Nature Communications, 7(1), 1–8. https://doi.org/10.1038/ncomms12452

Beirão, J., Purchase, C., Wringe, B., & Fleming, I. (2014). Wild Atlantic cod sperm motility is negatively affected by ovarian fluid of farmed females. Aquaculture Environment Interactions, 5(1), 61–70. https://doi.org/10.3354/aei00095

Bird, R. B., Hassager, R. C., & Armstrong, C. F. C. (2021). Kinetic Theory. Dynamics of Polymeric Liquids, 1(86), 204–207. https://doi.org/10.1017/S0022112078211081

Birkhead, T. R., & Pizzari, T. (2002). Postcopulatory sexual selection. Nature Reviews Genetics, 3(4), 262–273. https://doi.org/10.1038/NRG774

Böni, L., Fischer, P., Böcker, L., Kuster, S., & Rühs, P. A. (2016). Hagfish slime and mucin flow properties and their implications for defense. Scientific Reports, 6(1), 1–8. https://doi.org/10.1038/srep30371

Brokaw, C. J. (1965). Non-sinusoidal bending waves of sperm flagella. Journal of Experimental Biology, 43(1), 155–169. https://europepmc.org/article/med/5894323

Brokaw, C. J. (1966). 113-139 I I 3 With 2 plates and 14 text-figures. In J. Exp. Biol (Vol. 45).

Brokaw, C. J. (1983). Control Mechanisms in Sperm Flagella. In The Sperm Cell (pp. 291–303). Springer Netherlands. https://doi.org/10.1007/978-94-009-7675-7_53

Butts, I. A. E., Johnson, K., Wilson, C. C., & Pitcher, T. E. (2012). Ovarian fluid enhances sperm velocity based on relatedness in lake trout, Salvelinus namaycush. Theriogenology, 78(9), 2105-2109.e1. https://doi.org/10.1016/j.theriogenology.2012.06.031

Butts, I. A. E., Prokopchuk, G., Kašpar, V., Cosson, J., & Pitcher, T. E. (2017). Ovarian fluid impacts flagellar beating and biomechanical metrics of sperm between alternative reproductive tactics. Journal of Experimental Biology, 220(12), 2210–2217. https://doi.org/10.1242/jeb.154195

Cosson, J. (2015). Flagellar mechanics and sperm guidance. Bentham Science Publishers.

Cosson, J. (2021). A 40 years journey with fish spermatozoa as companions as I personally experienced it. Fish Physiology and Biochemistry, 47(3), 757–765. https://doi.org/10.1007/S10695-020-00882-

Cosson, J., Groison, A. L., Suquet, M., Fauvel, C., Dreanno, C., & Billard, R. (2008). Marine fish spermatozoa: Racing ephemeral swimmers. In Reproduction (Vol. 136, Issue 3, pp. 277–294). BioScientifica. https://doi.org/10.1530/REP-07-0522

Cosson, J., & Prokopchuk, G. (n.d.). Wave propagation in flagella. Retrieved September 9, 2020, from https://www.researchgate.net/publication/288260551

Cosson, M. P., Billard, R., Gatti, J. L., & Christen, R. (1985). Rapid and quantitative assessment of trout spermatozoa motility using stroboscopy. Aquaculture, 46(1), 71–75. https://doi.org/10.1016/0044-8486(85)90178-4

Cox, W. P., & Merz, E. H. (1958). Correlation of dynamic and steady flow viscosities. Journal of Polymer Science, 28(118), 619–622. https://doi.org/10.1002/POL.1958.1202811812

Dadras, H., Dzyuba, B., Cosson, J., Golpour, A., Siddique, M. A. M., & Linhart, O. (2017). Effect of water temperature on the physiology of fish spermatozoon function: a brief review. In Aquaculture Research (Vol. 48, Issue 3, pp. 729–740). Blackwell Publishing Ltd. https://doi.org/10.1111/are.13049

Dadras, H., Dzyuba, V., Cosson, J., Golpour, A., & Dzyuba, B. (2016). The in vitro effect of temperature on motility and antioxidant response of common carp Cyprinus carpio spermatozoa. Journal of Thermal Biology, 59, 64–68. https://doi.org/10.1016/j.jtherbio.2016.05.003

Dziewulska, K., Rzemieniecki, A., Czerniawski, R., & Domagała, J. (2011). Post-thawed motility and fertility from Atlantic salmon (Salmo salar L.) sperm frozen with four cryodiluents in straws or pellets. Theriogenology, 76(2), 300–311. https://doi.org/10.1016/j.theriogenology.2011.02.007

Dziewulska, K., Rzemieniecki, A., & Domagała, J. (2011). Sperm motility characteristics of wild Atlantic salmon (Salmo salar L.) and sea trout (Salmo trutta m. trutta L.) as a basis for milt selection. Journal of Applied Ichthyology, 27(4), 1047–1051. https://doi.org/10.1111/j.1439-0426.2011.01759.x

Eisenbach, M., & Giojalas, L. C. (2006). Sperm guidance in mammals — an unpaved road to the egg. Nature Reviews Molecular Cell Biology 2006 7:4, 7(4), 276–285. https://doi.org/10.1038/nrm1893

Elofsson, H., Mcallister, B. G., Kime, D. E., Mayer, I., & Borg, B. (2003). Long lasting stickleback sperm; is ovarian fluid a key to success in fresh water? Journal of Fish Biology, 63(1), 240–253. https://doi.org/10.1046/j.1095-8649.2003.00153.x

Engineering Properties of Foods. (2014). In Engineering Properties of Foods. CRC Press. https://doi.org/10.1201/9781420028805

Evans, J. P., & Lymbery, R. A. (2020). Sexual selection after gamete release in broadcast spawning invertebrates. Philosophical Transactions of the Royal Society B: Biological Sciences, 375(1813), 20200069. https://doi.org/10.1098/rstb.2020.0069

Evans, J. P., Rosengrave, P., Gasparini, C., & Gemmell, N. J. (2013). Delineating the roles of males and females in sperm competition. Proceedings of the Royal Society B: Biological Sciences, 280(1772). https://doi.org/10.1098/RSPB.2013.2047

Evans, J. P., & Sherman, C. D. H. (2013). Sexual Selection and the Evolution of Egg-Sperm Interactions in Broadcast-Spawning Invertebrates.

Fernández, J. L., Muriel, L., Rivero, M. T., Goyanes, V., Vazquez, R., & Alvarez, J. G. (2003). The Sperm Chromatin Dispersion Test: A Simple Method for the Determination of Sperm DNA Fragmentation. Journal of Andrology, 24(1), 59–66. https://doi.org/10.1002/J.1939-4640.2003.TB02641.X

Firman, R. C., Gasparini, C., Manier, M. K., & Pizzari, T. (2017). Postmating Female Control: 20 Years of Cryptic Female Choice. In Trends in Ecology and Evolution (Vol. 32, Issue 5, pp. 368– 382). Elsevier Ltd. https://doi.org/10.1016/j.tree.2017.02.010

Fitzpatrick, J. L., Bridge, C. D., & Snook, R. R. (2020). Repeated evidence that the accelerated evolution of sperm is associated with their fertilization function. Proceedings of the Royal Society B: Biological Sciences, 287(1932), 20201286. https://doi.org/10.1098/rspb.2020.1286

Fitzpatrick, J. L., & Lüpold, S. (2014). Sexual selection and the evolution of sperm quality. In Molecular Human Reproduction (Vol. 20, Issue 12, pp. 1180–1189). Oxford University Press. https://doi.org/10.1093/molehr/gau067

Fitzpatrick, J. L., Simmons, L. W., & Evans, J. P. (2012). Complex patterns of multivariate selection on the ejaculate of a broadcast spawning marine invertebrate. Evolution, 66(8), 2451–2460. https://doi.org/10.1111/j.1558-5646.2012.01627.x

Galvano, P. M., Johnson, K., Wilson, C. C., Pitcher, T. E., & Butts, I. A. E. (2013). Ovarian fluid influences sperm performance in lake trout, Salvelinus namaycush. Reproductive Biology, 13(2), 172–175. https://doi.org/10.1016/j.repbio.2013.02.001

Gasparini, C., Andreatta, G., & Pilastro, A. (2012). Ovarian fluid of receptive females enhances sperm velocity. Naturwissenschaften, 99(5), 417–420. https://doi.org/10.1007/s00114-012-0908-2

Gasparini, C., & Pilastro, A. (2011). Cryptic female preference for genetically unrelated males is mediated by ovarian fluid in the guppy. Proceedings of the Royal Society B: Biological Sciences, 278(1717), 2495–2501. https://doi.org/10.1098/rspb.2010.2369

Herschel, & H., W. (1926). Measurement of consistency of rubber-benzene solutions. Kolloid-Zeitschrift, 39, 291–298. https://ci.nii.ac.jp/naid/10014919340

Hirano, T., Morisawa, M., & Suzuki, K. (1978). Changes in plasma and coelomic fluid composition of the mature salmon (Oncorhynchus keta) during freshwater adaptation. Comparative Biochemistry and Physiology -- Part A: Physiology, 61(1), 5–8. https://doi.org/10.1016/0300-9629(78)90266-9

Holwill, M. E. J. (1977). Some Biophysical Aspects of Ciliary and Flagellar Motility. Advances in Microbial Physiology, 16(C), 1–48. https://doi.org/10.1016/S0065-2911(08)60046-6

Hussain, Y. H., Guasto, J. S., Zimmer, R. K., Stocker, R., & Riffell, J. A. (2016). Sperm chemotaxis promotes individual fertilization success in sea urchins. https://doi.org/10.1242/jeb.134924

Hyakutake, T., Sato, K., & Sugita, K. (2019). Study of bovine sperm motility in shear-thinning viscoelastic fluids. Journal of Biomechanics, 88, 130–137. https://doi.org/10.1016/j.jbiomech.2019.03.035

Immler, S., Hotzy, C., Alavioon, G., Petersson, E., & Arnqvist, G. (n.d.). Sperm variation within a single ejaculate affects offspring development in Atlantic salmon. https://doi.org/10.1098/rsbl.2013.1040

Ingermann, R. L., Bencic, D. C., & Gloud, J. G. (2001). Low seminal plasma buffering capacity corresponds to high pH sensitivity of sperm motility in salmonids. Fish Physiology and Biochemistry, 24(4), 299–307. https://doi.org/10.1023/A:1015037422720

Johnson, S. L., Borziak, K., Kleffmann, T., Rosengrave, P., Dorus, S., & Gemmell, N. J. (2020). Ovarian fluid proteome variation associates with sperm swimming speed in an externally fertilizing fish. Journal of Evolutionary Biology, 33(12), 1783–1794. https://doi.org/10.1111/jeb.13717

Kekalainen, J., & Evans, J. P. (2018). Gamete-mediated mate choice: Towards a more inclusive view of sexual selection. In Proceedings of the Royal Society B: Biological Sciences (Vol. 285, Issue 1883). Royal Society Publishing. https://doi.org/10.1098/rspb.2018.0836

Kelly, D. A., & Moore, B. C. (2016). The Morphological Diversity of Intromittent Organs. Integrative and Comparative Biology, 56(4), 630–634. https://doi.org/10.1093/ICB/ICW103

Kholodnyy, V., Gadêlha, H., Cosson, J., & Boryshpolets, S. (2019). How do freshwater fish sperm find the egg? The physicochemical factors guiding the gamete encounters of externally fertilizing freshwater fish. https://doi.org/10.1111/raq.12378

Lahnsteiner, F., Weismann, T., & Patzner, R. A. (n.d.). Composition of the ovarian fluid in 4 salmonid species: Oncorhynchus mykiss, Salmo trutta f lacustris, Saivelinus alpinus and Hucho hucho.

Lauga, E. (2007). Propulsion in a viscoelastic fluid. Physics of Fluids, 19(8), 83104. https://doi.org/10.1063/1.2751388

Lüpold, S., Manier, M. K., Berben, K. S., Smith, K. J., Daley, B. D., Buckley, S. H., Belote, J. M., & Pitnick, S. (2012). How multivariate ejaculate traits determine competitive fertilization success in drosophila melanogaster. Current Biology, 22(18), 1667–1672. https://doi.org/10.1016/j.cub.2012.06.059

Manier, M. K., Belote, J. M., Berben, K. S., Lüpold, S., Ala-Honkola, O., Collins, W. F., & Pitnick, S. (2013). Rapid Diversification Of Sperm Precedence Traits And Processes Among Three Sibling Drosophila Species. Evolution, 67(8), 2348–2362. https://doi.org/10.1111/evo.12117

Manier, M. K., Lüpold, S., Belote, J. M., Starmer, W. T., Berben, K. S., Ala-Honkola, O., Collins, W. F., & Pitnick, S. (2013). Postcopulatory Sexual Selection Generates Speciation Phenotypes in Drosophila. Current Biology, 23(19), 1853–1862. https://doi.org/10.1016/J.CUB.2013.07.086

Markovitz, H. (1981). Viscoelastic properties of polymers. Journal of Colloid and Interface Science, 84(2), 552. https://doi.org/10.1016/0021-9797(81)90247-2

Metz, E. C., Kane, R. E., Yanagimachi, H., & Palumbi, S. R. (2016). Fertilization Between Closely Related Sea Urchins Is Blocked by Incompatibilities During Sperm-Egg Attachment and Early Stages of Fusion. https://Doi.Org/10.2307/1542162, 187(1), p23–34. https://doi.org/10.2307/1542162

Oliver, M., & Evans, J. P. (2014). Chemically moderated gamete preferences predict offspring fitness in a broadcast spawning invertebrate. Proceedings of the Royal Society B: Biological Sciences, 281(1784). https://doi.org/10.1098/rspb.2014.0148

Palumbi, S. R. (1999). All males are not created equal: Fertility differences depend on gamete recognition polymorphisms in sea urchins. Proceedings of the National Academy of Sciences, 96(22), 12632–12637. https://doi.org/10.1073/PNAS.96.22.12632

Parker, G. A. (2020). Conceptual developments in sperm competition: a very brief synopsis. Philosophical Transactions of the Royal Society B: Biological Sciences, 375(1813), 20200061. https://doi.org/10.1098/rstb.2020.006.

Pitnick, S. S., Hosken, D. J., & Birkhead, T. R. (Eds.). (2008). Sperm biology: an evolutionary perspective. Academic press.

Poli, F., Immler, S., & Gasparini, C. (2019). Effects of ovarian fluid on sperm traits and its implications for cryptic female choice in zebrafish. Behavioral Ecology, 30(5), 1298–1305. https://doi.org/10.1093/beheco/arz077

Purchase, C. F., & Rooke, A. C. (2020). Freezing ovarian fluid does not alter how it affects fish sperm swimming performance: creating a cryptic female choice “spice rack” for use in split-ejaculate experimentation. https://doi.org/10.1111/jfb.14263

Ramón, M., Jiménez-Rabadán, P., García-Álvarez, O., Maroto-Morales, A., Soler, A. J., Fernández-Santos, M. R., Pérez-Guzmán, M. D., & Garde, J. J. (2014). Understanding sperm heterogeneity: Biological and practical implications. Reproduction in Domestic Animals, 49(4), 30–36. https://doi.org/10.1111/RDA.12404

Rooke, A. C., Palm-Flawd, B., Purchase, C. F., & Cooke, S. (2019). The impact of a changing winter climate on the hatch phenology of one of North America’s largest Atlantic salmon populations. 7. https://doi.org/10.1093/conphys/coz015

Rosengrave, P., Taylor, H., Montgomerie, R., Metcalf, V., McBride, K., & Gemmell, N. J. (2009). Chemical composition of seminal and ovarian fluids of chinook salmon (Oncorhynchus tshawytscha) and their effects on sperm motility traits. Comparative Biochemistry and Physiology - A Molecular and Integrative Physiology, 152(1), 123–129. https://doi.org/10.1016/j.cbpa.2008.09.009

Schoff, C. K., & Kamarchik, P. (2005). Rheology and Rheological Measurements. In Kirk-Othmer Encyclopedia of Chemical Technology. John Wiley & Sons, Inc. https://doi.org/10.1002/0471238961.1808051519030815.a01.pub2

Simmons, L. W. (2005). The Evolution of Polyandry: Sperm Competition, Sperm Selection, and Offspring Viability. Annual Review of Ecology, Evolution, and Systematics, 36(1), 125–146. https://doi.org/10.1146/annurev.ecolsys.36.102403.112501

Sloan, N. S., & Simmons, L. W. (2019). The evolution of female genitalia. Journal of Evolutionary Biology, 32(9), 882–899. https://doi.org/10.1111/JEB.13503

Storm, C., Pastore, J. J., MacKintosh, F. C., Lubensky, T. C., & Janmey, P. A. (2005). Nonlinear elasticity in biological gels. Nature, 435(7039), 191–194. https://doi.org/10.1038/nature03521

Suarez, S. S., & Pacey, A. A. (2006). Sperm transport in the female reproductive tract. In Human Reproduction Update (Vol. 12, Issue 1, pp. 23–37). Oxford Academic. https://doi.org/10.1093/humupd/dmi047

Swanson, W. J., & Vacquier, V. D. (1997). The abalone egg vitelline envelope receptor for sperm lysin is a giant multivalent molecule. Proceedings of the National Academy of Sciences, 94(13), 6724– 6729. https://doi.org/10.1073/PNAS.94.13.6724

Taylor, M. L., Price, T. A. R., & Wedell, N. (2014). Polyandry in nature: a global analysis. Trends in Ecology & Evolution, 29(7), 376–383. https://doi.org/10.1016/J.TREE.2014.04.005

Turner, E., & Montgomerie, R. (2002). Ovarian fluid enhances sperm movement in Arctic charr. Journal of Fish Biology, 60(6), 1570–1579. https://doi.org/10.1006/jfbi.2002.2018

Weir, L. K., Breau, C., Hutchings, J. A., & Cunjak, R. A. (2010). Multiple paternity and variance in male fertilization success within Atlantic salmon Salmo salar redds in a naturally spawning population. Journal of Fish Biology, 77(3), 479–493. https://doi.org/10.1111/j.1095-8649.2010.02690.x

Wojtczak, M., Dietrich, G. J., Słowińska, M., Dobosz, S., Kuźmiński, H., & Ciereszko, A. (2007). Ovarian fluid pH enhances motility parameters of rainbow trout (Oncorhynchus mykiss) spermatozoa. Aquaculture, 270(1–4), 259–264. https://doi.org/10.1016/j.aquaculture.2007.03.010

Wolfner, M. F. (2011). Precious Essences: Female Secretions Promote Sperm Storage in Drosophila. PLOS Biology, 9(11), e1001191. https://doi.org/10.1371/JOURNAL.PBIO.1001191

Yanagimachi, R., Cherr, G. N., Pillai, M. C., & Baldwin, J. D. (1992). Factors Controlling Sperm Entry into the Micropyles of Salmonid and Herring Eggs. (fish/sperm/egg/micropyle/fertilization). Development, Growth and Differentiation, 34(4), 447–461. https://doi.org/10.1111/j.1440-169X.1992.00447.x

Yanagimachi, R., Harumi, T., Matsubara, H., Yan, W., Yuan, S., Hirohashi, N., Iida, T., Yamaha, E., Arai, K., Matsubara, T., Andoh, T., Vines, C., & Cherr, G. N. (2017). Chemical and physical guidance of fish spermatozoa into the egg through the micropyle†,‡. Biology of Reproduction, 96(4), 780–799. https://doi.org/10.1093/BIOLRE/IOX015

Yeates, S. E., Diamond, S. E., Einum, S., Emerson, B. C., Holt, W. v., & Gage, M. J. G. (2013). Cryptic choice of conspecific sperm controlled by the impact of ovarian fluid on sperm swimming behavior. Evolution, 67(12), 3523–3536. https://doi.org/10.1111/evo.12208

Zadmajid, V., Myers, J. N., Sørensen, R., Anthony, I., & Butts, E. (2019). Ovarian fluid and its impacts on spermatozoa performance in fish: A review. https://doi.org/10.1016/j.theriogenology.2019.03.021

