## supplementary material for "Frequency-dependent viscosity of salmon ovarian fluid has biophysical implications for sperm-egg interactions"

***Preliminary assessment of the ovarian fluid rheology and preservation method***

After filtration and pH, volume, conductivity and density measurements were taken (see materials and methods section), each batch of ovarian fluid was divided in three separate falcon vials containing equal volumes that were then stored at - 80, -20, and 4 °C respectively. A preliminary rheological analysis, using a portion of these samples (N= 5) was conducted to assess the best preservation method. Being that the techniques used to assess the viscoelastic profiles of the samples were particularly time consuming (1 or 2 samples per day maximum), we wanted to avoid any bacterial degradation in the 4°C-stored samples that would eventually affect the fluid’s structure. Nevertheless, we also wanted to ensure that the freezing thermal treatments would not affect the polymeric structure of the fluid. No observable hysteresis was found between the three thermal treatments when subjected to the experiments illustrated. Therefore, we opted for the samples stored at -20 °C for optimal processing through the duration of the experiments, and to these all the results are referred.

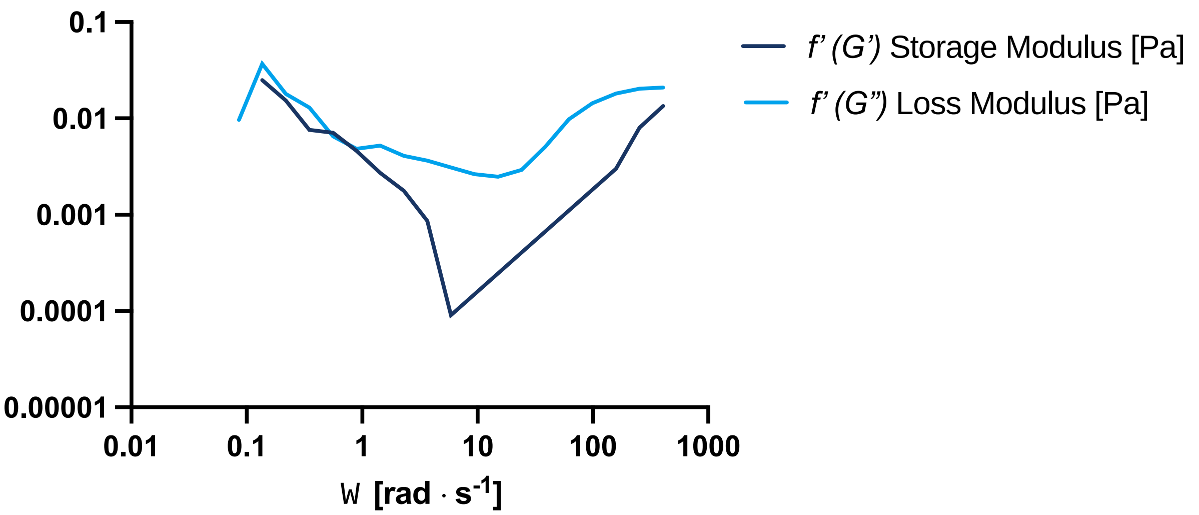


*Figure 1. First derivative (f’) of the Storage modulus (G') and loss modulus (G'') of Atlantic salmon ovarian fluids (n= 11), to describe the relation between the viscous and elastic components of the fluid at increasing angular frequencies (0 <Ω< 500 rad· $s^{-1}$). Data are presented as means.*

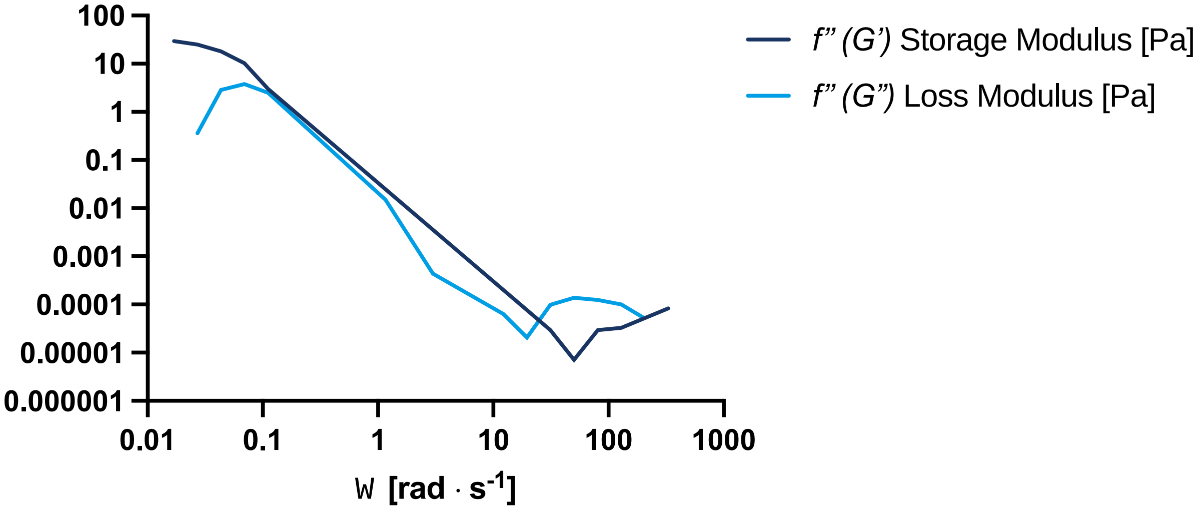


*Figure 2. Second derivative (f”) of the Storage modulus (G') and loss modulus (G'') of Atlantic salmon ovarian fluids (n= 11), to describe the relation between the viscous and elastic components of the fluid at increasing angular frequencies (0 <Ω< 500 rad· $s^{-1}$). Data are presented as means.*
